# Puberty Blocker and Aging Impact on Testicular Cell States and Function

**DOI:** 10.1101/2024.03.23.586441

**Authors:** Varshini Murugesh, Megan Ritting, Salem Salem, Syed Mohammed Musheer Aalam, Joaquin Garcia, Asma J Chattha, Yulian Zhao, David JHF Knapp, Guruprasad Kalthur, Candace F Granberg, Nagarajan Kannan

**Affiliations:** Department of Laboratory Medicine and Pathology, Mayo Clinic, Rochester, MN, USA; Department of Pediatrics, Mayo Clinic, Rochester, MN, USA; Division of Reproductive Endocrinology and Infertility, Department of Obstetrics and Gynecology, Mayo Clinic, Rochester, MN, USA; Institut de Recherche en Immunologie et Cancérologie, and Département de Pathologie et Biologie Cellulaire, Université de Montréal, Montreal, QC, Canada; Division of Reproductive Biology, Department of Reproductive Science, Kasturba Medical College, Manipal, Manipal Academy of Higher Education, Manipal, India; Department of Urology, Mayo Clinic, Rochester, MN, USA; Mayo Clinic Comprehensive Cancer Center, Mayo Clinic, Rochester, MN, USA; Center for Regenerative Biotherapeutics, Mayo Clinic, Rochester, MN, USA

**Keywords:** Spermatogonial stem cell, Sertoli cell, Leydig cell, Testicular development, Puberty, Puberty Blocker, GnRH, GnRH analog, Transgender Medicine

## Abstract

Spermatogonial stem cell (SSC) acquisition of meiotogenetic state during puberty to produce genetically diverse gametes is blocked by drugs collectively referred as ‘puberty blocker’ (PB). Investigating the impact of PB on juvenile SSC state and function is challenging due to limited tissue access and clinical data. Herein, we report largest clinically annotated juvenile testicular biorepository with all children with gender dysphoria on chronic PB treatment highlighting shift in pediatric patient demography in US. At the tissue level, we report mild-to-severe sex gland atrophy in PB treated children. We developed most extensive integrated single-cell RNA dataset to date (>100K single cells; 25 patients), merging both public and novel (52 month PB-treated) datasets, alongside innovative computational approach tailed for germ cells and evaluated the impact of PB and aging on SSC. We report novel constitutional ranges for each testicular cell type across the entire age spectrum, distinct effects of treatments on prepubertal vs adult SSC, presence of spermatogenic epithelial cells exhibiting post-meiotic-state, irrespective of age, puberty status, or PB treatment. Further, we defined distinct effects of PB and aging on testicular cell lineage composition, and SSC meiotogenetic state and function. Using single cell data from prepubertal and young adult, we were able to accurately predict sexual maturity based both on overall cell type proportions, as well as on gene expression patterns within each major cell type. Applying these models to a PB-treated patient that they appeared pre-pubertal across the entire tissue. This combined with the noted gland atrophy and abnormalities from the histology data raise a potential concern regarding the complete ’reversibility’ and reproductive fitness of SSC. The biorepository, data, and research approach presented in this study provide unique opportunity to explore the impact of PB on testicular reproductive health.

## Introduction

Spermatogonial stem cells (SSC) undergo a developmental progression originating from early embryonic stem cells, transitioning through primordial germ cells, and eventually differentiating into spermatogonia (SPG), marking a crucial trajectory in reproductive cell development. SSC represents a rare (<0.05 %) subset of SPG that compensates for cell loss and maintains the undifferentiated reservoir through periodic self-renewal divisions while giving rise to transit amplifying SPG poised for differentiation (*1, 2*). The differentiating spermatogonia undergo rapid and stochastic mitotic divisions before entering meiosis as spermatocytes (SPC), leading to the formation of haploid spermatids and ultimately, functional spermatozoa (*3*).

The transition from childhood to sexual maturity (i.e. puberty) involves substantial changes in testicular physiology enabling somatic cell maturation and initiating spermatogenesis following SSC’s initiation of meiotogenetic function (*4, 5*). A rhesus monkey study in 1980 showed that in females ovulatory cycles could be reversibly induced in prepubertal females through periodic gonadotropin-releasing hormone (GnRH) infusion followed by reversal to an immature state upon regimen discontinuation (*6*), but similar studies in male are not reported. Nevertheless, these findings heralded the notion that neither adenohypophysial nor gonadal competence imposes limitations on puberty initiation (*6*), and suggested that the meiotogenetic state, in both male and female, is controlled by hypothalamus-pituitary-gonadal (HPG)-axis, through neurobiological mechanisms that first delay and then initiate the onset of puberty (*7*). The purportedly ‘reversible’ meiotogenetic state controlled by HPG-axis produce genetically diverse functional spermatozoa is highly conserved in mammals (*8, 9*). Understanding these mechanisms is essential for designing safe interventions and predicting long-term outcomes, yet current knowledge in human, particularly regarding the influence of drugs targeting HPG-axis in the SSC transition, remains limited (*10*).

The initiation of meiotogenesis, intricately linked with sexual maturation, is suggested to mark the onset of haploid cell production in the testis and dependent on crucial brain signal, GnRH. Manipulating GnRH levels through injection has led to a surge in gonadotropin levels in patients with puberty disorders, contributing to the development of GnRH-related testosterone (T) suppressants (*11, 12*). Pulsatile gonadotropin signals are initially observed at-birth and are subsequently reinitiated during the onset of puberty. The sexual developmental clock, in conjunction with permissive signals related to somatic growth, energy balance, and season, regulates the activation of GnRH neurons at the onset of meiotogenesis (*13*). This clock rhythmically controls GnRH neuron activity, transitioning from basal, non-pulsatile stimulation of pituitary gonadotropin release in early prepuberty to elevated pulsatile stimulation during the night in REM cycles in late prepuberty, and to regularized elevated pulsatile stimulation throughout adulthood (*12, 14, 15*). The role of hypothalamic KNDy neurons, presumed regulators of GnRH neurons in the sexual developmental clock, remains incompletely understood (*16*). Although the physiological control of this clock is not fully elucidated, it is apparent that the target cells in testis remain sensitive to the signals associated with this clock.

Puberty blockers (PB) are drugs that inhibit the re-initiation of pulsatile GnRH-1 decapeptide release from hypothalamic GnRH neurons thereby delaying the development of secondary sexual characteristics in adolescents. Most prominent PB namely GnRH analogs (GnRHa), initially developed to treat cancers, have found applications in treating paraphilic disorders (*17–19*), delaying sexual maturation in children with precocious puberty (*20, 21*), tertiary hypogonadism (*22*) and children with gender dysphoria (GD) (*23, 24*). Among GD patients, GnRHa treatment prevented pubertal development (*25*). Nevertheless, the consequences of exposure to PB on juvenile testicular development and reproductive fitness of SSC are poorly understood. A previous study in adult rat with short-term GnRH antagonist treatment suggested reversible effects on the gonadal function and fertility (*26*). However, a 1993 study on prepubertal bulls showed GnRHa treatment decreased LH pulse frequency, had no effect on basal LH, but increased T concentrations (*27*) and do not address the reproductive fitness following the treatment. To the best of our knowledge, no rigorous study has been reported on extended puberty blockade in pediatric populations and its long-term consequences on reproductive fitness.

Despite limited specimen availability, single cell RNA sequencing technology has significantly advanced our understanding of postnatal human testicular development providing a high-resolution transcriptomic map at the single-cell level (*10, 28–32*). These investigations have revealed distinct cell populations within the testicular microenvironment, shedding light on the heterogeneity and functional diversity of testicular cells. Key cell types identified include Sertoli cells, crucial for supporting spermatogonial development, Leydig cells responsible for T production, and spermatogonia, the germ cells that initiate spermatogenesis. Additionally, these studies have unveiled unique subpopulations and cellular states, contributing to a more nuanced comprehension of testicular biology. How these cell types change during aging and enable meitogenetic transition of SSC remains unexplored.

Herein, we analyzed testicular biorepository data from pediatric patients with and without chronic PB exposure (*33, 34*). We provide unprecedented histological evidence revealing detrimental pediatric testicular sex gland responses to PB. With the help of the first known testicular single-cell map of a 14-year-old patient with over 52 months of constant PB treatment reported herein and public data, we developed an integrated single-cell molecular analysis approach and define effects of PB and aging on testicular composition, SSC state, and meiotogenetic function during aging. Finally, employing machine learning approach, we developed a model using single cell data to predict personalized meiotogenetic fitness.

## Results

### Widespread Puberty Blocker Use Among Children with Gender Dysphoria Undergoing Fertility Preservation Surgery

Using pediatric testicular biorepository protocol which enrolls patients between 0-17 years of age and diagnosed with a fertility-threatening condition and/or are scheduled to undergo gonadotoxic treatment (*33, 34*) (see Methods; **Table 1** and **Figure 1A**), we have collected living testicular specimens for clinical and research applications from 92 individuals. Of these, 5 withdrew from the study, leaving 87 patients for further analysis. Among these, 55 (63 %) were oncology patients, 11 (12 %) had hereditary diseases, and 16 (18 %) were diagnosed with gender dysphoria (GD). All 16 GD patients identified as transgender female. The average age at the time of gender transition and fertility preservation (FP) surgery is 8.1 (age range = 2—15; std deviation = 4.6) and 12.5-years old (age range = 10—16; std deviation = 1.8), respectively. The average age of PB initiation is 12.1-years old (age range = 10—16.4; std deviation = 1.83). Remarkably, 100 % of GD patients were under PB treatment including 9 patients (56 %) at the time of FP surgery, highlighting the widespread nature of PB intervention in this demographic (**Table 1**). Although sperm collection was suggested, all GD patients opted for FP surgery due to reluctance or inability to ejaculate. Two out of 9 PB-treated patients exhibited abnormalities: one had bilateral abnormal testicles with a lack of complete tunica albuginea, while another had a right testis that was not easily palpable. The remaining patients displayed scrotum symmetry with bilaterally palpable testes.

**Figure 1:**
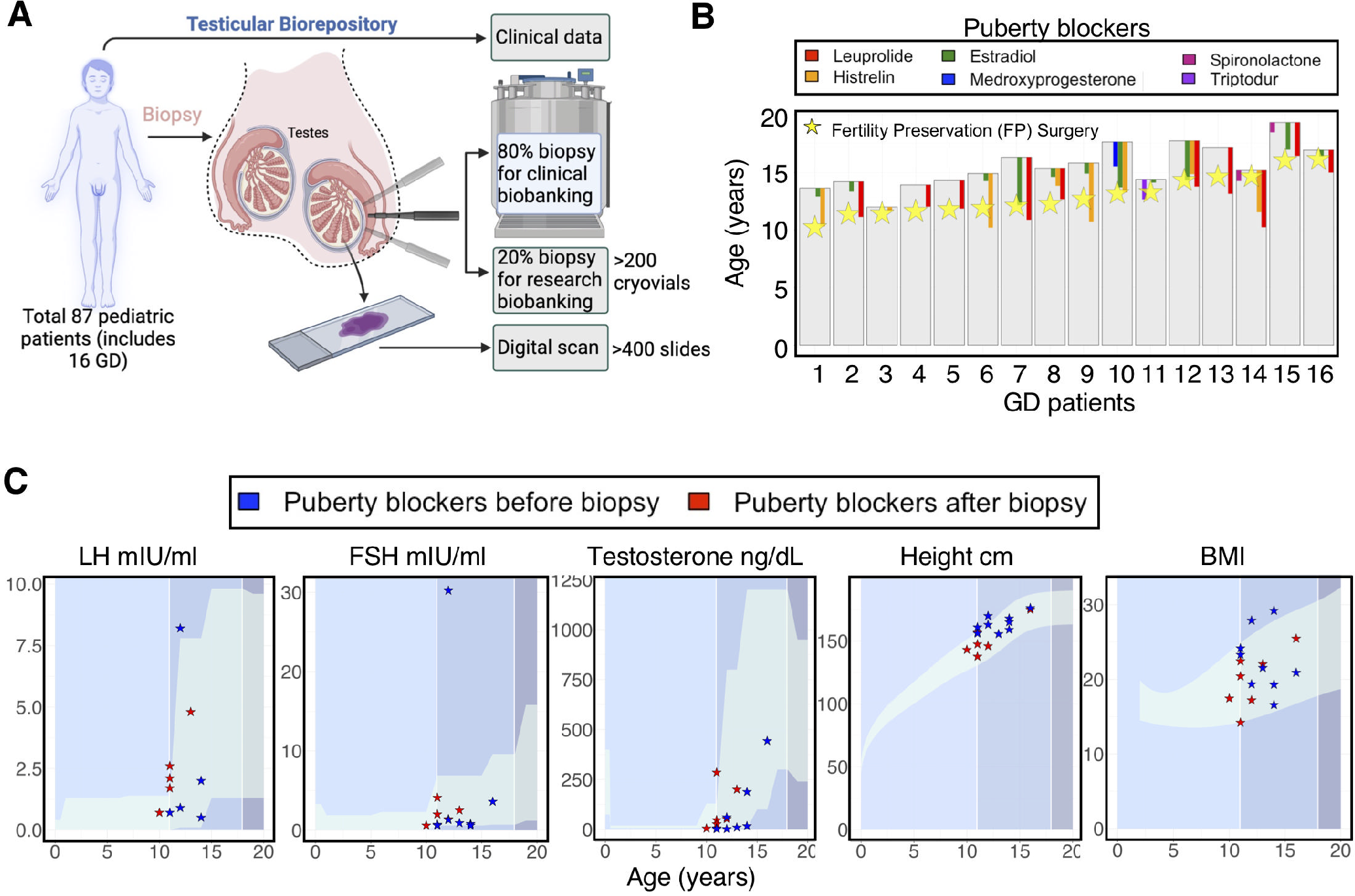
*Overview of pediatric testicular biorepository and clinical data for patients undergoing puberty blocker (PB) treatment.* A) Schematic representation of pediatric testicular biorepository. **B)** Clinical management of 16 GD patients showing PB treatment timeframes and age at fertility preservation (FP) surgery; **C)** Endocrine (LH, FSH and T) profile, height, and body-mass-index (BMI) of GD patients. Clinical laboratory reference ranges across age for the parameters are highlighted.

**Table 1:** Characteristics of donors and specimen of the Mayo Clinic’s pediatric testicular biobank for fertility preservation. All 16 pediatric patients with gender dysphoria received PB treatment, with 56 % at the time of surgery, shows shift in patient demography.

|  | All Patients | Non-GD patients |  |  |  | GD patients |  |
| --- | --- | --- | --- | --- | --- | --- | --- |
| Characteristic | 0-17 Years<br>N = 87 | 0-10 Years<br>N = 53 | 11-14 Years<br>N = 15 | 15-17 Years<br>N = 3 | 0-10 Years<br>N = 1 | 11-14 Years<br>N = 13 | 15-17 Years<br>N = 2 |
| **Race — no. (%)** |  |  |  |  |  |  |  |
| White | 77 (88.5%) | 49 (92.47%) | 13 (86.67%) | 3 (100%) | 1 (100%) | 10 (76.92%) | 1 (50%) |
| African American | 5 (5.7%) | 2 (3.77%) | 0 (0%) | 0 (0%) | 0 (0%) | 2 (15.38%) | 1 (50%) |
| White/ Hispanic or Latino | 2 (2.3%) | 1 (1.88%) | 0 (0%) | 0 (0%) | 0 (0%) | 1 (7.7%) | 0 (0%) |
| Asian | 1 (1.2%) | 1 (1.88%) | 0 (0%) | 0 (0%) | 0 (0%) | 0 (0%) | 0 (0%) |
| Other | 2 (2.3%) | 0 (0%) | 2 (13.33%) | 0 (0%) | 0 (0%) | 0 (0%) | 0 (0%) |
| **BMI — no. (%)** |  |  |  |  |  |  |  |
| <16 | 20 (23%) | 14 (26.41%) | 4 (26.67%) | 1 (33.33%) | 0 (0%) | 1 (7.7%) | 0 (0%) |
| 16-25 | 58 (66.7%) | 35 (66.05%) | 10 (66.66%) | 1 (33.33%) | 1 (100%) | 10 (76.92%) | 1 (50%) |
| 25-30 | 6 (6.8%) | 3 (5.66%) | 0 (0%) | 0 (0%) | 0 (0%) | 2 (15.38%) | 1 (50%) |
| >30 | 2 (2.3%) | 0 (0%) | 1 (6.67%) | 1 (33.33%) | 0 (0%) | 0 (0%) | 0 (0%) |
| Unknown | 1 (1.2%) | 1 (1.88%) | 0 (0%) | 0 (0%) | 0 (0%) | 0 (0%) | 0 (0%) |
| **Gender at Surgery — no. (%)** |  |  |  |  |  |  |  |
| Male | 69 (79.3%) | 52 (98.12%) | 14 (93.33%) | 3 (100%) | - | - | - |
| Female | 16 (18.4%) | - | - | - | 1 (100%) | 13 (100%) | 2 (100%) |
| Not disclosed | 2 (2.3%) | 1 (1.88%) | 1 (6.67%) | 0 (0%) | 0 (0%) | 0 (0%) | 0 (0%) |
| **Genetic History — no. (%)** |  |  |  |  |  |  |  |
| Yes | 28 (32.18%) | 22 (41.5%) | 4 (26.67%) | 0 (0%) | 0 (0%) | 1 (7.7%) | 0 (0%) |
| No | 59 (67.82%) | 31 (58.5%) | 11 (73.33%) | 3 (100%) | 1 (100%) | 12 (92.3%) | 2 (100%) |
| **Surgical History — no. (%)** |  |  |  |  |  |  |  |
| Yes | 70 (80.45%) | 48 (90.56%) | 13 (86.67%) | 2 (66.67%) | 0 (0%) | 7 (53.85%) | 0 (0%) |
| No | 17 (19.55%) | 5 (9.44%) | 2 (13.33%) | 1 (33.33%) | 1 (100%) | 6 (46.15%) | 2 (100%) |
| **Preexisting Medical Condition — no. (%)** |  |  |  |  |  |  |  |
| Oncological | 55 (63.17%) | 42 (79.26%) | 11 (73.33%) | 2 (66.67%) | 0 (0%) | 0 (0%) | 0 (0%) |
| Hereditary disease | 11 (12.6%) | 8 (15.09%) | 3 (20%) | 0 (0%) | 0 (0%) | 0 (0%) | 0 (0%) |
| Kleinfelter Syndrome | 1 (1.2%) | 0 (0%) | 0 (0%) | 1 (33.33%) | 0 (0%) | 0 (0%) | 0 (0%) |
| Torsion | 1 (1.2%) | 0 (0%) | 1 (6.67%) | 0 (0%) | 0 (0%) | 0 (0%) | 0 (0%) |
| Gender Dysphoria | 16 (18.33 %) | 0 (0%) | 0 (0%) | 0 (0%) | 1 (100%) | 13 (100%) | 2 (100%) |
| None | 1 (1.2%) | 1 (1.88%) | 0 (0%) | 0 (0%) | 0 (0%) | 0 (0%) | 0 (0%) |
| Other | 2 (2.3%) | 2 (3.77%) | 0 (0%) | 0 (0%) | 0 (0%) | 0 (0%) | 0 (0%) |
| **History of Mental Trauma — no. (%)** |  |  |  |  |  |  |  |
| Yes | 2 (2.3%) | 1 (1.88%) | 0 (0%) | 0 (0%) | 0 (0%) | 1 (7.7%) | 0 (0%) |
| No | 85 (97.7%) | 52 (98.12%) | 15 (100%) | 3 (100%) | 1 (100%) | 12 (92.3%) | 2 (100%) |
| **Suicidal Tendency — no. (%)** |  |  |  |  |  |  |  |
| Yes | 3 (3.44%) | 0 (0%) | 0 (0%) | 0 (0%) | 0 (0%) | 2 (15.38%) | 1 (50%) |
| No | 84 (96.56%) | 53 (100%) | 15 (100%) | 3 (100%) | 1 (100%) | 10 (84.62%) | 1 (50%) |
| **Tanner Staging — no. (%)** |  |  |  |  |  |  |  |
| 1 | 48 (55.17%) | 39 (73.6%) | 6 (39.99%) | 0 (0%) | 1 (100%) | 0 (0%) | 0 (0%) |
| 2 | 10 (11.49%) | 1 (1.88%) | 2 (13.33%) | 0 (0%) | 0 (0%) | 2 (15.38%) | 0 (0%) |
| 3 | 7 (8.06%) | 0 (0%) | 4 (26.67%) | 0 (0%) | 0 (0%) | 7 (53.85%) | 0 (0%) |
| 4 | 4 (4.6%) | 0 (0%) | 1 (6.67%) | 1 (33.33%) | 0 (0%) | 3 (23.07%) | 2 (100%) |
| 5 | 3 (3.44%) | 0 (0%) | 1 (6.67%) | 2 (66.67%) | 0 (0%) | 0 | 0 (0%) |
| Unknown | 15 (17.24%) | 13 (24.52%) | 1 (6.67%) | 0 (0%) | 0 (0%) | 1 (7.7%) | 0 (0%) |
| Research Tissue Weight in g — mean (range) | 0.057<br>(0.003-0.519) | 0.035<br>(0.0039-0.468) | 0.101<br>(0.024-0.519) | 0.098<br>(0.06-0.129) | 0.111 | 0.078<br>(0.022-0.243) | 0.147<br>(0.065-0.229) |
| **Sperm — no. (%)** |  |  |  |  |  |  |  |
| Present | 12 (13.8%) | 0 (0%) | 5 (33.33%) | 1 (33.33%) | 1 (100%) | 4 (30.76%) | 1 (50%) |
| Absent | 63 (72.41%) | 47 (88.67%) | 5 (33.33%) | 1 (33.33%) | 0 (0%) | 9 (69.24%) | 1 (50%) |
| NA | 12 (13.79%) | 6 (11.33%) | 5 (33.33%) | 1 (33.33%) | 0 (0%) | 0 (0%) | 0 (0%) |
| Number of Plasma Vials — mean±SD | 0.30±0.84 | 0.352±0.86 | 0.533±1.18 | 0±0 | 0±0 | 0±0 | 0±0 |
| Number of Patient Vials — mean±SD | 6.4±1.43 | 5.826±1.04 | 7.38±1.5 | 8±2 | 8±0 | 7.46±1.71 | 7±0 |
| Number of Research Vials — mean±SD | 2.6±0.92 | 2.36±0.69 | 3.38±1.12 | 3.667±1.52 | 4±0 | 2.76±1.42 | 3±0 |
| **Puberty Blocker Treated — no. (%)** |  |  |  |  |  |  |  |
| Yes | 16 (18.33%) | 0 (0%) | 0 (0%) | 0 (0%) | 1 (100%) | 13 (100%) | 2 (100%) |
| No | 71 (81.67%) | 53 (100%) | 15 (100%) | 3 (100%) | 0 (0%) | 0 (0%) | 0 (0%) |
| **Puberty Blocker Treatment Initiated — no. (%)** |  |  |  |  |  |  |  |
| Before FP Surgery | 9 (10.34%) | 0 (0%) | 0 (0%) | 0 (0%) | 0 (0%) | 8 (61.54%) | 1 (50%) |
| After FP Surgery | 78 (89.66%) | 53 (100%) | 15 (100%) | 3 (100%) | 1 (100%) | 5 (38.46%) | 1 (50%) |

PB, including leuprolide, histrelin, triptorelin, estradiol, medroxyprogesterone, and spironolactone, were administered with exposure durations ranging from 3 to 52 months at the time of surgery is depicted in **Figure 1B**. Behavioral assessments indicated higher (Fischer’s exact test; p = 0.0057) suicidal tendencies among patients with GD compared to those without (**Table 1**). GD patients exposed to PB exhibited levels of FSH, LH, T, height, and BMI within the constitutional range, though their levels were generally closer to the lower limits for FSH and T, suggesting lack of complete suppression of the HPG-axis by these treatments (**Figure 1C**).

### Degenerated Sex Gland Development Associated with Puberty Blocker Treatment

Typically, healthy testes exhibit uniform seminiferous tubule (sex gland) development. However, previous findings have highlighted a prevalent occurrence of mixed patterns, especially within the hypospermatogenesis classification in men (*35*). The extent of heterogeneous intra-sex gland spermatogenesis in biopsies from pubertal tissues— whether exposed to PB or not—remains inadequately understood.

For the sex gland study, we digitally scanned 400+ Hematoxylin & Eosin-stained donor testicular biopsy sections available in Mayo Clinic tissue registry. We analyzed testicular specimen with and without PB exposure. The histology results showed abnormal testicular development in PB treated compared to non-treated patients (**Figure 2A-B**).

**Figure 2:**
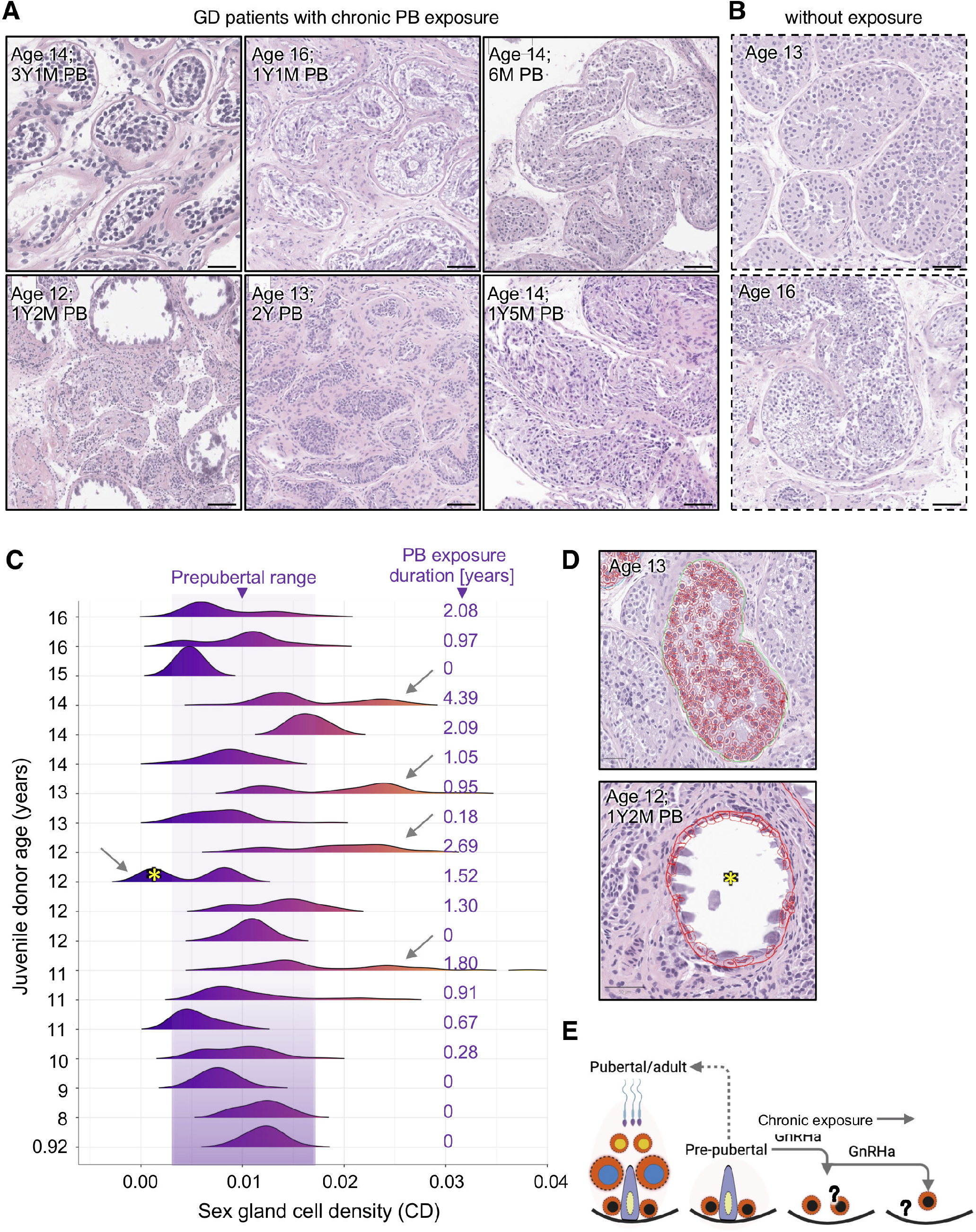
***Puberty blocker and juvenile aging on sex gland development.*** Representative images of Hematoxylin and Eosin-stained sections of testicular tissue biopsied from the testis from GD patients **(A)** with and **(B)** without PB exposure. **C)** Ridgeline plot showing sex gland cell density (CD) across 19 samples (3 non-GD and 16 GD patients) including 9 treated with PB (as shown in Figure 1B). A Linear regression analysis of CD across patients to test association between CD and age was performed. P value was calculated using Wald test. Arrows highlight abnormal CD distribution. **D)** QuPath-based computerized scoring method for digitally scanned stained cells within sex gland cross-sectional area. Representative images of normal (top) and fully atrophied sex gland (bottom). Star represents fully atrophied sex glands. **E)** A proposed model of chronic effects of PB treatment on male sex glands.

To gain deeper insights, we identified 19 donors aged 0.92 to 16 years, all without a history of cancer. We analyzed ≤ 50 individual sex glands per donor for both cross-sectional area and cell count using QuPath software (see Methods). The sex gland cross-sectional cell densities (CD) were calculated. The data revealed that prepubertal sex glands were generally smaller and with greater CD than pubertal. Increase in distinct bimodal distribution of CD in patients especially in PB-treated indicated two distinct response states of sex glands (**Figure 2C**). Analysis of bimodal peaks suggested that some sex glands maintained prepubertal-like CD whereas others abnormally densely packed. Testicular development in various juvenile patients exhibited heterogeneous responses to PB, indicating inter-patient variability. For example, sex glands of a 12-year-old patient treated with leuprolide and estradiol (E2) for a period of 14 months had 59 % of sex glands fully atrophied with appearance of microlithiasis (**Figure 2C-D**).

Linear regression analysis of CD across donors showed highly significant (Wald test; p-value = 3.81 x 10^-10^) association with age in patients without PB treatment, and an insignificant (p-value = 0.872) association with age in patients with PB treatment. A prior study noted variations in the maturation rates of Sertoli cells across distinct sex glands in pubertal tissue (*10*). Our tissue-level analysis quantifies the heterogeneous development of sex glands, highlighting the substantial impact of PB treatment on germ epithelial homeostasis within these structures (**Figure 2E**).

### Single-Cell Analysis Reveals the Impact of Puberty Blocker and Aging on Testicular Lineage Composition

Given the complexity of the developing juvenile sex glands and the physical interaction of functionally distinct cell types in the testis, we used scRNA-seq approach to dissect the molecular pathologies of prolonged puberty blocker treatment on testicular cell states. We obtained fresh left testicular tissue from a 14-year-old patient with over 52 months of PB treatment. The tissue was enzymatically dissociated to isolate single cells, and scRNA-seq library were generated and sequenced as previously described (*28*). Our analysis included publicly available testicular single-cell RNA data across postnatal ages and data from Guo et al. of two adults undergoing gender confirmation surgery treated with either E2 alone or with T antagonist spironolactone therapy (referred to hereafter as T-suppressant), totaling 130,100 cells, including 11,199 from our juvenile PB-treated GD patient representing largest single cell analysis of testes (see Methods)(*10, 28, 30, 32*).

Employing 146 established markers and a uniform manifold approximation and projection (UMAP) plot for dimensionality reduction, we identified 24 clusters corresponding to known testicular cell populations (**Figure 3A-D; Supplementary Figure S1A-C** and **Supplementary Figure S2A-D**). Testicular UMAPs of different developmental stages and drug exposures are presented in **Figure 3A**. Analysis of spermatogenic, Sertoli and Leydig lineage composition, showed abnormal testicular cell development in PB-treated and T-suppressant-treated patients compared to untreated controls (**Figure 3B**).

**Figure 3:**
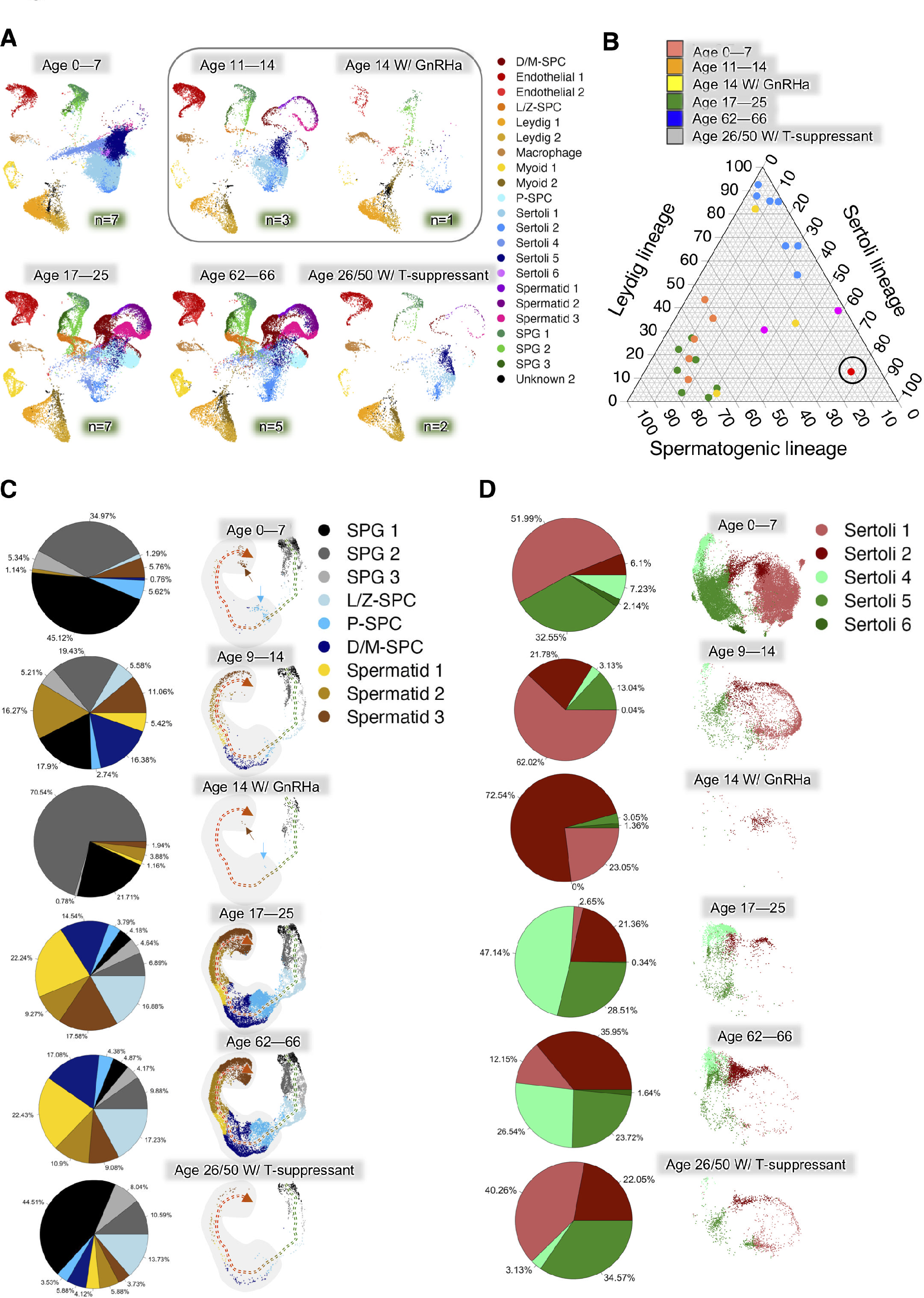
***Impact of puberty blocker and aging on testicular cell composition.* A)** UMAP projection of all identified cell types split between age group and treatment received. **B)** Ternary plot of spermatogenic, Sertoli and Leydig cell lineages of testicular tissue of prepubertal (0-7-year old), prepubertal receiving PB (14-year old), pubertal, young adult (11-14-year old), young adult receiving T-suppressant (26 and 50-year old) and old adult (62-66-year old). **C)** Comparative assessment of changes in relative proportion of spermatogenic cell linages with normal development (n = 22), T- suppressant (n = 2), and PB (n = 1) treatment. Dashed arrows represent trajectory of differentiation. Solid arrows highlight post-meiotic stages of development. **D)** Comparative assessment of changes in relative proportion of Sertoli cell lineages with normal development (n = 22), T-suppressant (n = 2), and PB (n = 1) treatment.

Based on the cell distributions from normal samples over time, we established constitutional ranges for each across ontogeny using the convex hull method (see Methods) (**Figure 4A-B** and **Supplementary Figure S3**).

**Figure 4:**
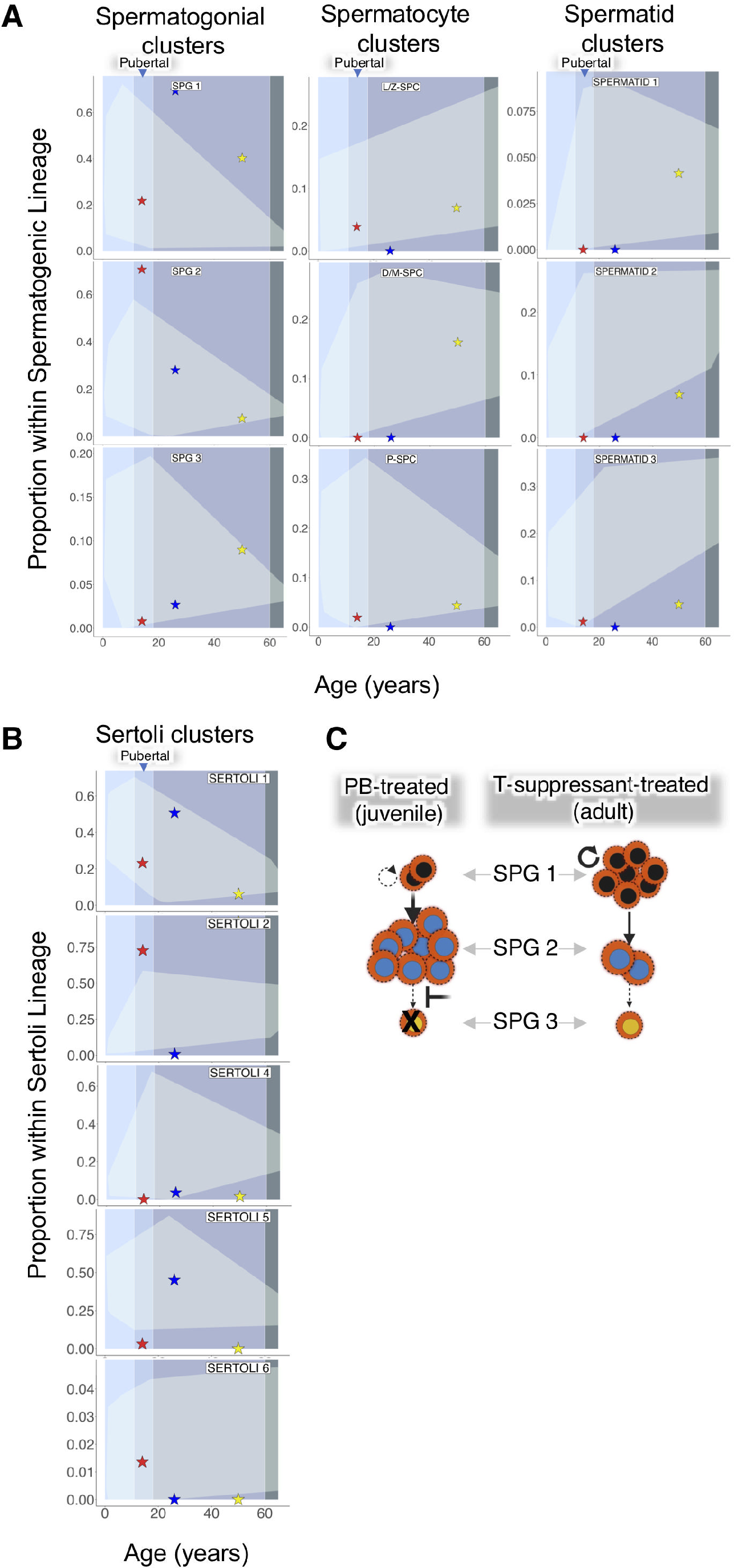
***Impact of puberty blocker and aging on spermatogonial and Sertoli cell development.* A)** Trends in the relative proportion of spermatogenic cell lineages with normal development. The red star represents the PB-treated patient, whereas blue and yellow stars respectively represent the 26-year-old and 50-year-old T-suppressant- treated patients. **B)** Trends in the relative proportion of Sertoli cell lineages with normal development are highlighted in light blue. The star color remains the same. **C)** A model depicting the effects of PB exposure in juvenile patients, illustrating abnormally elevated SPG 2 cell type and differentiation block, while T-suppressant exposure in adults leads to the depletion of SPG 1 cell type.

We observed distinct lineage contributions between prepubertal and adult patients. Spermatogenic epithelial cell type distribution during puberty clearly marked progression from prepubertal towards adult like pattern (**Figure 3B**). Notably, prepubertal, PB-treated juvenile and T-suppressant-treated adults exhibited SSC differentiation patterns, deviating from pubertal and adult testes, respectively (**Figure 3B-C**). Further analysis revealed a developmental block at the spermatogonial state, predominantly observed in PB-treated patients. Additionally, abnormal proportions of specific spermatogenic and Sertoli cell states were evident in the PB-treated patient compared to expected normal ranges (**Figure 3A-D**; **Figure 4A-C** and **Supplementary Figure S3**).

In the PB-treated patient, a proportion of cells within the spermatogenic lineage indicated a developmental block at the spermatogonial state (> 90 % of cells) (**Figure 3B**), and pathologically higher levels of Sertoli 4 and lower levels of Sertoli 1/5 states (**Figure 3C**). Additionally, abnormal frequencies of specific spermatogenic and Sertoli cell types were evident in the PB-treated patient compared to patients with normal pubertal development (**Figure 3C-D** and **Figure 4A-B**). PB-treated juvenile patient had abnormally higher representation of SPG 2 and in comparison, T-suppressant-treated adult patients had abnormally higher representation of SPG 1 suggesting that juvenile and adult testes may experience distinct developmental blockades to HPG-axis manipulation (**Figure 4C**).

### Post-meiotogenic-state, Although Rare, is Evident in Newborns and Prepubertal Individuals

Meiotogenesis is generally assumed to be absent in normal pre-pubertal children due to lack of sperm development. Remarkably, we observed spermatid-like cells across all juvenile stages, including newborns. The proportions of pre-meiotic (SPG 1/2/3, L/Z-SPC) versus post-meiotic (P/D/M-SPC, Spermatid 1/2/3) spermatogenic epithelial lineage reached up to 12 % of the total spermatogenic epithelium in prepubertal stages. In fact, PB-treated juvenile patient exhibited 7 % spermatids within spermatogenic epithelial lineage (**Figure 3C**; **Figure 4A** and **Supplementary Figure S3**).

We revealed a striking molecular clustering of pre-pubertal spermatid-like cells, challenging conventional assumptions about their developmental trajectory post-pubertal onset (**Figure 3C**). Furthermore, our analysis of significantly differentially expressed genes in pre-pubertal spermatid 2 and 3 compared to their adult counterparts suggests a potentially novel induction pathway independent of puberty (**Supplementary Table S1** and **Supplementary Table S2**). This pathway leads to likely incomplete differentiation, resulting in the development of spermatid-like cells without subsequent sperm maturation and warrants investigation.

### Puberty-associated Gene (PAG) Analysis Identifies Distinct Cell-specific Changes

Genes related to puberty have been pinpointed using genome-wide association studies (GWAS), which examine the entire genome for associations with puberty-related traits. Additionally, pathogenic gene variants known to affect puberty have been identified (*36, 37*). However, we still lack a clear understanding of how these genes are expressed in different types of cells in the testicles at different ages. We explored the expression patterns of these puberty-associated genes (PAGs) across different testicular cell types and examined whether they exhibited any age-specific changes (**Figure 5A-B** and **Supplementary Figure S3**).

**Figure 5:**
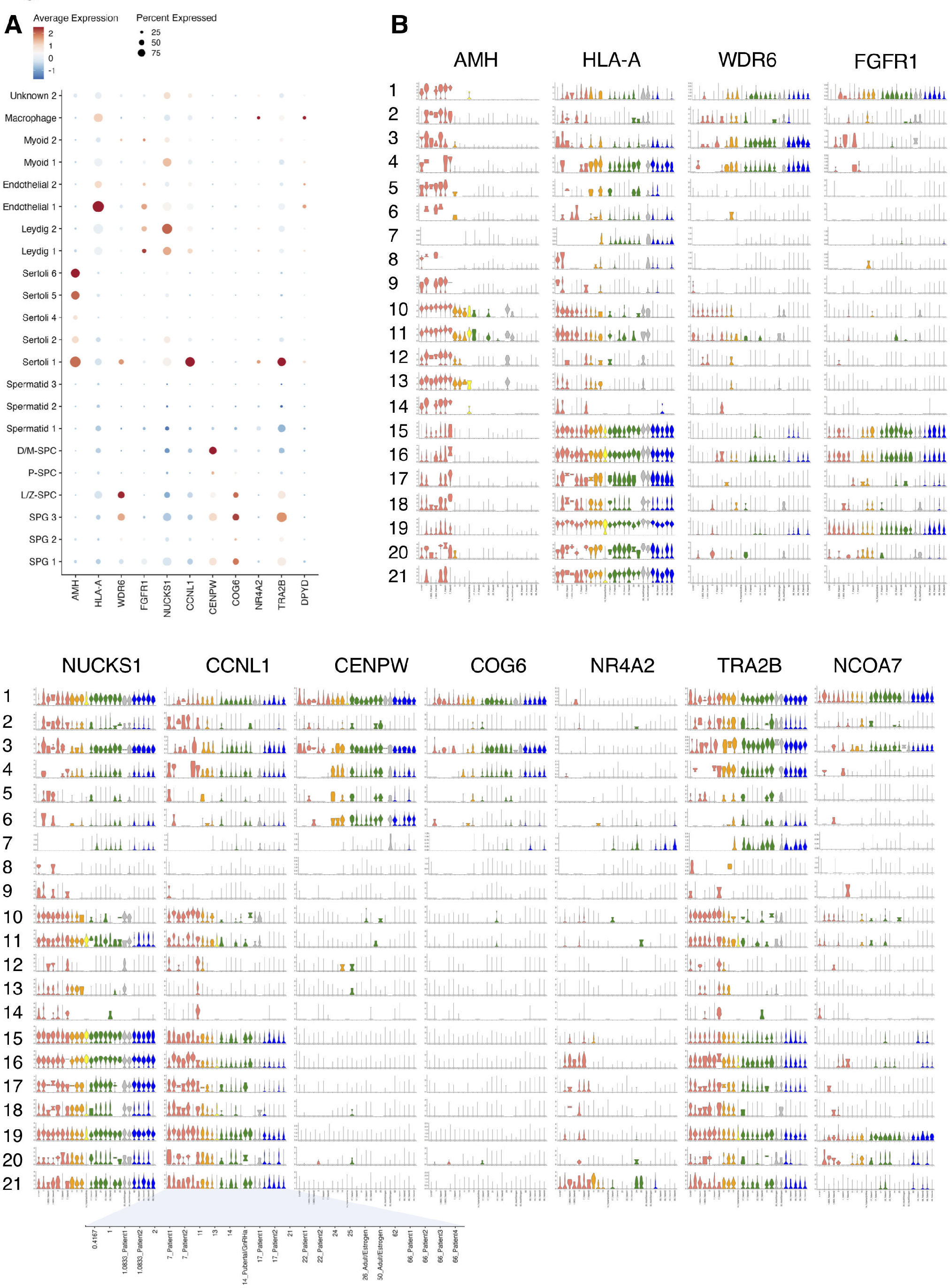
***Impact of puberty blocker and aging on expression of puberty-associated genes (PAGs).* A)** Dot plot depicting relative expression of select PAGs across all cell types. **B)** Violin plots representing the expression patterns of select PAGs across cell types and age. Cell types are denoted by numbers viz., 1=SPG 1; 2=SPG 2; 3=SPG 3; 4=L/Z-SPC; 5=P-SPC; 6=D/M-SPC; 7=Spermatid 1; 8=Spermatid 2; 9=Spermatid 3; 10=Sertoli 1; 11=Sertoli 2; 12=Sertoli 4; 13=Sertoli 5; 14=Sertoli 6; 15=Leydig 1; 16=Leydig 2; 17=Myoid 1; 18=Myoid 2; 19=Endothelial 1; 20=Endothelial 2; and 21=Macrophage.

The normalized scRNAseq data were merged using the RNA assay method (see Methods). We detected 174 PAGs in our datasets, showing lineage-specific expression: 173 in the spermatogenic lineage, 170 in the Sertoli lineage, 164 in the Leydig lineage, 161 in the myoid lineage, 166 in the endothelial lineage, and 155 in macrophage cells (**Supplementary Table S3**). We illustrate complex cell-specific, age-related alterations in PAG expression. The data also delineates distinct impacts of treatments with PB during juvenile and T-suppressant in adults (**Figure 5A**).

Subsequently, we examined their expression profiles across chronological age, revealing previously undiscovered developmental trends for these PAGs. Firstly, Anti-Müllerian hormone (AMH), pivotal for GnRH neuron development (*38*), and Sertoli cell maturation from fetal stages to puberty (*39, 40*), exhibited high expression across all testicular cell types, contrary to previous assumptions of exclusive Sertoli cell expression. Notably, all testicular cell types downregulated AMH expression in adults, with pubertal suppression following this order: SPG 1/Sertoli 5 > Sertoli 4/2/1 > Leydig 1/2 > Endothelial 1/2 (**Figure 5B** and **Supplementary Figure S5**).

Other noteworthy age-related patterns include: CCNL1 expression until late prepuberty in Leydig 1, NUCKS3 expression in spermatid 3, and NR4A2 expression in myoid 1. Additionally, TRA2B expression persisted until late puberty in myoid 2, WDR6 in Sertoli 1, and HLA-A in Sertoli 5 (**Figure 5B** and **Supplementary Figure S5**). We observed elevated DPYD expression in tissue-resident macrophages during late puberty, a phenomenon mitigated by PB and T-suppressants (**Figure 5A**). Distinct expression patterns of PAGs were noted for prepubertal post-meiotic cells, although their significance remains unclear.

Furthermore, the endothelial 1 cluster exhibited ubiquitous expression of FGFR1 and anti-inflammatory NCOA7 across ages, yet PB treatment suppressed their expression in juveniles, unlike T-suppressant treatment in adults (**Figure 5B**). This suggests potential age-related variations in testicular endothelial cell responses to these agents. Our findings lend support to the notion that PB treatment may promote juvenile testicular atrophy partly through testicular endothelial dysfunction.

### Changes in Transcriptomic States of Primitive Spermatogonial Cells (SPG 1 and 2)

Furthermore, SPG 1 and 2 cell transcriptomes are significantly altered in the PB-treated patient. SPG 2 cell transcriptomes of the PB patient had greater number of differentially regulated genes when compared to SPG 1, as observed in comparison to pubertal stage and T-suppressant-treated adults (**Figure 6A** and **6B**).

**Figure 6:**
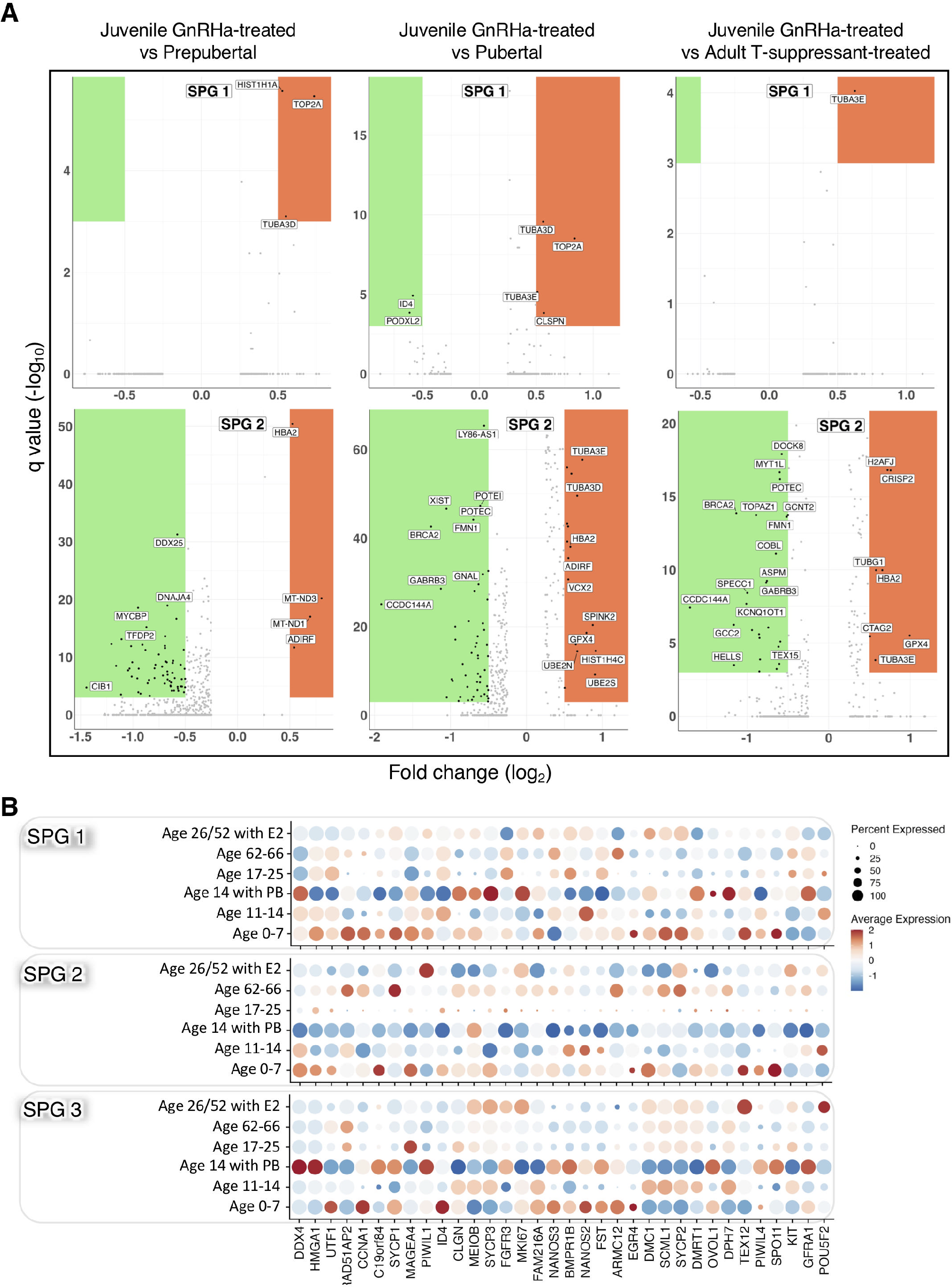
***Puberty blocker affects spermatogonial stem cell state.* A)** Volcano plot showing differential downregulated genes in green and upregulated genes in orange of the PB-treated patient SPG cells compared to those of prepubertal (n = 7), pubertal (n = 3), and T-suppressant-treated (n = 2). Q-value threshold used was 0.001. SPG 3 was excluded from analysis since the PB-treated patient did not have enough cells to perform differential expression analysis. Y-axis denotes -log10 of q-values whereas x-axis denotes log2 of fold change values. Volcano plots were generated using *Seurat*, *qvalue*, and *ggplot2*. **B)** Dot plot depicting relative expression of standard spermatogenic lineage cell markers from existing literature between developmental categories and treatment received, if applicable.

In **Figure 6A**, the gene expression patterns shed light on key molecular changes associated with PB and T-suppressant treatments in SPG 1-state. Notably, prepubertal boys under PB treatment exhibit elevated expression of chromatin organization genes (HIST1H1A and TOP2A) and a microtubule formation gene (TUBA3D) compared to juvenile testicular tissue. This suggests potential regulatory shifts in chromatin organization and microtubule dynamics during PB intervention. In contrast, pubertal SPG 1 transcripts display significant upregulation of chromatin organization genes (TUBA3D, TOP2A, TUBA3B) and the cell cycle checkpoint regulator (CLSPN), coupled with downregulation of DNA binding inhibitory protein (ID4) and cell surface transmembrane protein (PODXL2) when compared to PB-treated prepubertal boys. This complex gene expression profile implies intricate molecular alterations associated with the onset of puberty. Conversely, the comparison between SPG 1 of adult men and T-suppressant- treated adult testicular cells reveals significant upregulation of TUBA3E, indicating potential hormonal influences on microtubule dynamics in adult testicular tissue. These findings provide valuable insights into the dynamic gene expression landscape associated with puberty, PB treatment, and hormonal exposure in germ cells of testis.

The observed gene expression changes in SPG 2 cells following PB treatment provide valuable insights into the molecular alterations associated with this intervention. Downregulation of genes involved in RNA secondary structure maintenance (DDX25), chaperone binding activity (DNAJA4), cMYC binding protein (MYCBP), transcription factor DP family member (TFDP2), and calcium ion binding (C1B1) suggests potential impacts on spermatogenesis and steroidogenesis. In the comparison between juvenile PB-treated and pubertal spermatogonial-2 cells, the downregulation of genes like LY86- A51, XIST, POTEC, POTE1, FMN1, BRCA2, GNAL, GABRB3, CCDC144A, and the upregulation of genes including TUB3E, TUBA3D, HBA2, ADIRF, VCX2, SPINK2, GPX4, HIST1H4C, UBE2N, and UBE2S highlight complex regulatory changes potentially influencing germ cell development in testis. Additionally, in the comparison between juvenile PB-treated and adult T-suppressant-treated spermatogonial-2 cells, observed gene expression changes point towards alterations in GTPase interaction (DOCK8), microvilli assembly (COBL), mitotic spindle regulation (ASPM), epigenetic modification (KCNQ10T1), and double-strand DNA repair (TEX15), alongside upregulation of genes related to histone regulation (H2AFJ), cell-cell adhesion (CRISP2), autoimmunogenic tumor antigen expression (CTAG2), and other key factors (**Figure 6A** and **6B**). Similarly, **Supplementary Figure S6** depicts genetic variances between PB-treated and pubertal Leydig, Sertoli, Myoid, and endothelial cell types. These findings suggest an impact of PB and T-suppressant treatments on the molecular landscape of SPG 2 cells, providing a basis for further exploration of their roles in germ cell development in testis.

### Testicular Targets of Sexual Developmental Clock Predict Spermatogenic Fitness

The initiation of meiotogenesis, marking the onset of haploid cell production in the testis, is intricately linked with sexual maturation. Previous studies suggest that while sex glands during prepuberty are not immature or incapable of gametogenesis, the crucial brain signal, GnRH, is lacking. Manipulating GnRH levels through injection has led to a surge in gonadotropin levels in patients with puberty disorders, contributing to the development of GnRH-related puberty blockers (*11, 12*). Pulsatile gonadotropin signals are initially observed at-birth and are subsequently reinitiated during the onset of puberty. The sexual developmental clock, in conjunction with permissive signals related to somatic growth, energy balance, and season, regulates the activation of GnRH neurons at the onset of meiotogenesis (*13*). This clock rhythmically controls GnRH neuron activity, transitioning from basal, non-pulsatile stimulation of pituitary gonadotropin release in early prepuberty to elevated pulsatile stimulation during the night in REM cycles in late prepuberty, and to regularized elevated pulsatile stimulation throughout adulthood (*12, 14, 15*). The role of hypothalamic KNDy neurons, presumed regulators of GnRH neurons in the sexual developmental clock, remains incompletely understood (*16*). Although the physiological control of this clock is not fully elucidated, it is evident that the target cells remain sensitive to the signals associated with this clock. Consequently, we sought to gain further insights into changes related to spermatogenic maturity and its modulation by age and the use of PB and T suppressants in patients.

As a first step, we trained a logistic regression model to classify individuals based on their proportion of SPG, Sertoli, and Leydig cells into adult or pre-pubertal classes (**Figure 7A**). Five prepubertal patients ages 13 months (n = 2), 2 years old (n = 1), and 7 years old (n = 2) and 5 young adult patients ages 21, 22 (n = 2), 24, and 25 were used to train this model. Within these individuals, classification accuracy and the area-under-the receiver operator curve (AUC) is provided in **Supplementary Table S4**. These values suggest that the model is generally accurate, suggesting that cell type composition can predict an individual’s spermatogenic maturity. It should be noted, however, that the low number of data points represents a major limitation for generalizability. To this point, one 5-month neonatal patient was misclassified as mature. None-the-less, applying this predictor to other samples yielded interesting information. For one, certain old adult patients were classified as prepubertal suggesting a loss of competency in spermatogenic lineage cells during aging. Finally, the PB-treated patient and the 26-year-old T-suppressant-treated patient were classified as prepubertal, whereas the 50-year-old T-suppressant-treated patient was classified as mature (**Figure 7B**). This suggests the idea that the age at which a drug is administered can affect a patient’s testicular niche.

**Figure 7:**
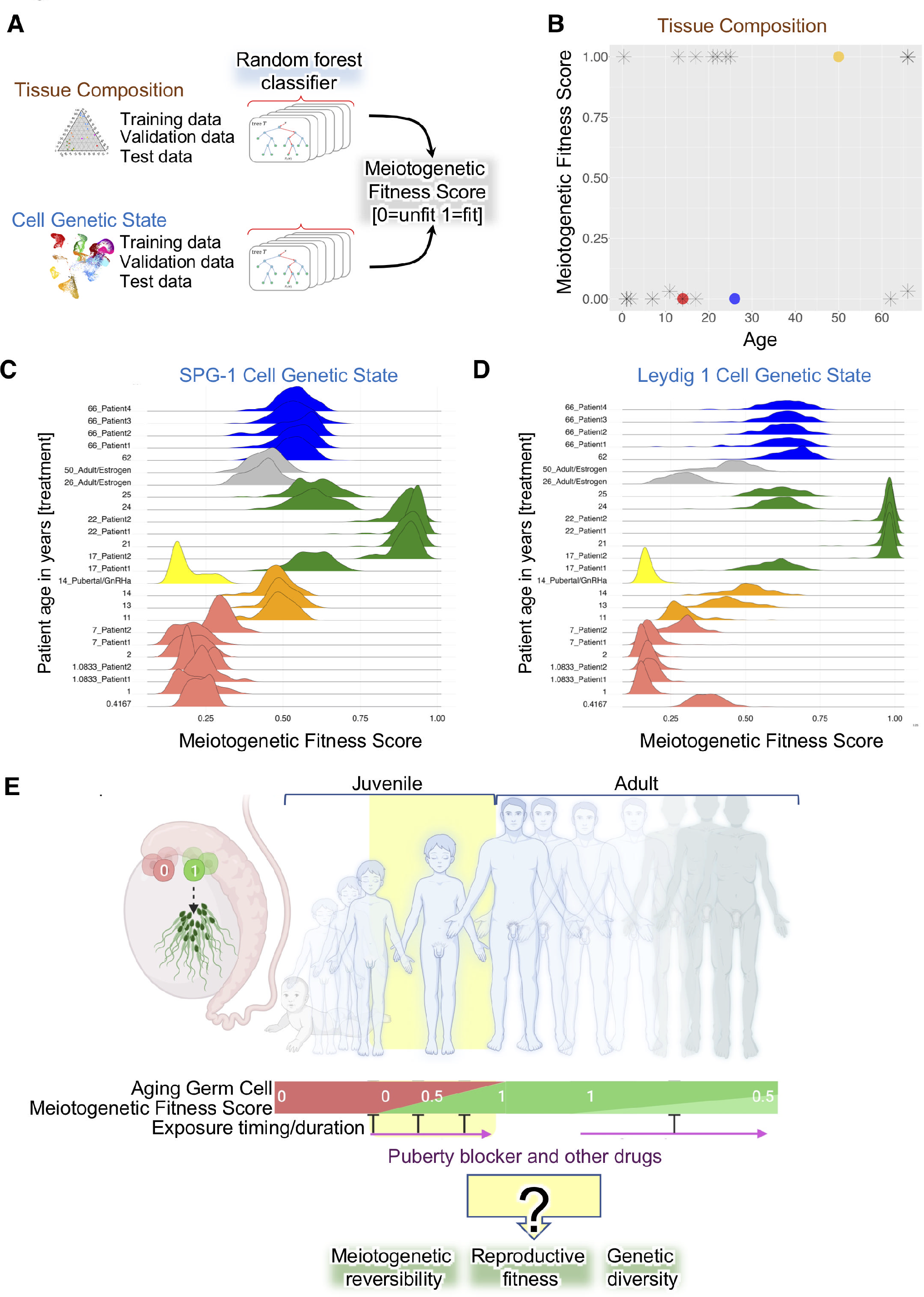
*Machine learning models predict meiotogenetic fitness at single cell resolution.* A) Schematic representation of random forest machine learning approach. **B)** Logistic regression model prediction scores with PB-treated patient in red, 26-year-old T-suppressant-treated adult in blue, and 50-year-old T-suppressant-treated adult in yellow. Distribution of prediction scores for **(C)** SPG 1 and **(D)** cells split by patient with color representing category of development. **D)** Random forest classifier table showing default thresholds selected by F1 score for each cluster, validation specificity, and validation sensitivity. **E)** Illustration demonstrating the utility of machine learning models in analyzing testicular single cells across age to study the effects of puberty blocker treatments on meiotogenetic reversibility, reproductive fitness, and the impact on healthy genetic diversity through the normal spermatogenesis process.

We next wanted to see whether in addition to differences in the cell type compositions, whether there were also changes in gene expression within these that reflected spermatogenic maturity. To do so, we trained a separate Random Forest classifier based on each cell type. Interestingly, strong predictive power was observed for all tested cell types. This suggests that shifts towards spermatogenic maturity likely result in broad microenvironmental changes and gene expression changes throughout the organ (**Figure 7C**). We next applied these models more broadly across the datasets to investigate how age and PB treatment affect cell type maturity. Pubertal samples tended to have intermediate prediction scores, suggesting that they may be in a transition state (**Figure 7D** and **Supplementary Figure S7**). Scores from prepubertal individuals were uniformly low as expected, but interestingly, some adult individuals also had more intermediate scores suggesting a degree of variability in the maturity-associated gene programs. Consistent with the logistic regression model based on cell type proportions, older individuals also tended to have lower scores suggesting an intermediate competence (**Figure 7D** and **Supplementary Figure S7**). Adults with T-suppressant treatment similarly had intermediate scores, generally even lower than older adults, also suggesting lower spermatogenic maturity. As expected, the GnRHa patient showed a range of per-cell scores similar to prepubertal across all examined cell populations (**Figure 7D** and **Supplementary Figure S7**).

Finally, we extracted predictive genes for meiotogenesis from the trained models, distinguishing between spermatogenic and non-spermatogenic lineage cellular states within the testis using a random forest classifier. To elucidate specific maturity-related programs, we compared the top contributing genes across different cell types (see matrix). The top 10 predictive genes across cell types are presented here (**Supplementary Figure S8**):

SPG 1: *SPACA3, SPATA18, SPANXC, ACR, PSORS1C1, FAM71F1, C7orf61, SPATA12, EQTN,* and *RPGRIP1*.

SPG 2: *GLIPR1L1, NPIPB6, LINC00964, TMEM225, SPANXC, GOLGA8S, EDRF1-AS1, TEX22, CCNA1,* and *LINC00226*.

Leydig 1: *ARMC12, TTC29, PCP2, CABYR, ERICH2, CMTM2, TUBA3C, SPATA18, ZPBP,* and *SPATA4*.

Leydig 2: *APLNR, DNAJC5B, COX7B2, ADAD1, FAM209B, ADCY4, SPERT, SPAG17, TUBA3E,* and *LINC00226*.

Sertoli 2: *SSX3, DHRS9, POTEI, ISL1, AREG, SLN, PXDNL, LIN28B, COBL,* and *RBM24*.

Sertoli 4: *KCNN3, ADAM32, NRGN, TMC7, IGF2BP2, LIN28B, C14orf39, NEFM, PCSK2,* and *IGSF10*.

Sertoli 5: *ELSPBP1, UNC13C, DPEP1, MYT1L, IGF1, DACH1, ITGA1, LIN28B, FILIP1,* and *PREX2*.

Several predictive genes have known role in spermatogenesis, greatly varied across cell types, suggesting different responses of these cells in response to sexual development (**Supplementary Figure S9**). The full list of predictive genes and their scaled prediction scores across these cell types are presented in **Supplementary Table S5**. Collectively, our approach provides new insights into predictive genes associated with spermatogenic maturity.

## Discussion

The pristine testicular tissues of a healthy juvenile populace are reserved primarily for fertility preservation and consequently, their availability to the scientific community is profoundly restricted. Therefore, our clinically annotated juvenile testicular specimens with and without PB exposure provide unique opportunity to understand puberty and PB response and conduct translational research. This is particularly significant given the escalating concern surrounding the decreasing age of pubertal onset in males, alongside the increasing utilization of transgender medicines in clinical practice, despite limited data on the long-term consequences of PB exposure on testicular reproductive health (*41*).

In males, the puberty-associated development of spermatogonial stem cells (SSC) undergoes dynamic cellular and molecular changes orchestrated through Sertoli and Leydig cells, influenced by intricate signals under the control of GnRH (*42*). GnRH analogs, versatile drugs, are increasingly used in clinics to treat Juvenile-GD by blocking the development of sexual characteristics, impeding both primary (reproductive) and secondary (non-reproductive) organ development in children of both sexes with GD (*20, 21*). GD patients who seek fertility preservation surgery face a major conundrum due to the absence of current procedures guaranteeing fertility (*43, 44*). This is because the generation of gametes from human SSC that have not naturally transitioned during puberty to a meiotogenetic state have been unsuccessful (*45, 46*). Prepubertal human testicular tissue/SSC has also failed to generate gametes in vitro or when xenografted in animals (*47, 48*). Additionally, the actions of GnRH and GnRHa may be influenced by genetics and lifestyle, making it challenging to predict individual responses to their SSC pubertal transition. The impact of chronic GnRHa, more potent and with longer half-life than natural GnRH, on juvenile testicular development and SSC meiotogenetic fitness, is poorly understood (*49*). The choice, dosage, and duration of GnRHa drugs for long-term gender dysphoria treatment during juvenescence and their effects on SSC are unclear, and there are currently no established clinical guidelines for discontinuation. The Mayo Clinic Pediatric Testicular Biobank for Fertility Preservation is a comprehensive resource addressing knowledge gaps. It is the largest pediatric (age 0-17 years) testicular biobank, encompassing GD children with and without PB treatment. The biobank includes well-annotated clinical data, reproductive health data, digitized slides of testicular tissues, and cryospecimens suitable for single-cell omics analysis (*34*).

Single-cell transcriptomic atlases have revealed spermatogonial stem cell (SSC) differentiation and key regulators in various species, including mice (*50–52*), rats (*53*), monkeys (*29, 54*), and humans (*10, 32, 55, 56*). However, studies in humans often focus on orchidectomy samples from deceased individuals of the same age, lacking investigation into juvenile aging and puberty-associated changes (*29, 57–60*). To ensure comprehensive coverage, our study utilized compatible public and patient data, encompassing twenty-five distinct males aged 1-66 years, a 14-year-old patient continuously treated with GnRHa since the age of 9, and two adult patients with gender-affirming T-suppressant therapy. This approach resulted in a single-cell map seamlessly integrating pubertal processes with the aging context. Notably, our study provides the first atlas of testicular cells of a child with prolonged GnRHa treatment.

Furthermore, we developed a novel single cell-based framework that determines constitutional ranges of proportionality of lineage-specific cell types which are predictive of changes associated with puberty and PB treatment. Moreover, recognizing the complexity and dynamic nature of pubertal SSC state we have developed logistical regression and random forest-based machine learning model to measure reproductive fitness of SSC. Albeit with limited patients, the single cell data used for training and validation suggest this method could very well be used to understand juvenescence and drugs promoting and preventing SSC meiotogenetic fitness associated with reproductive fitness and gamete genetic diversity. Leveraging these models, we unearthed unique predictive gene signatures defining prepubertal (potential) versus adult (kinetic) states of SSC pertaining to meiotogenetic function. The diverse methods described here help systematically probe the wealth of single cell testicular data, unveiling the intricacies behind the SSC development throughout PB and aging treatment and study associated changes in SSC meitogenetic fitness for personalized transgender medicine (**Figure 7E**).

Our intriguing finding of prepubertal spermatids prompts fundamental questions: At what stage of development the sex glands acquire the capability to generate sperm? These inquiries are not unfounded; rather, they compel us to advance Harris’s argument that prepubertal sex glands may not be entirely immature (*8*), and sparse meiotogenic occurrences may be a normal aspect of the developmental process. Indeed, a study has reported that approximately 8% of sex glands in a 1 ½-year-old child with Hurler syndrome (*61*). Regardless, our findings suggest that a rare population of SPG may undergo spontaneous differentiation independent of the sexual developmental clock associated with puberty. Notably, the present single cell RNA technology does not capture mature spermatozoa, limiting our identification to spermatogenic differentiation up to spermatid 3 (elongated). Further investigation is warranted.

In examining the mechanisms driving pubertal changes, previous studies have pinpointed specific genes and identified modifications in signal such as AMH, androgen signaling (*62–65*). Our investigation into these genes, along with the assessment of AMH, revealed some unexpected discoveries. Notably, our analysis uncovered minimal overlap in the regulation of genes within spermatogonial stem cells (SSC), specifically SPG 1 and 2, when comparing juvenile-GnRHa to prepubertal, pubertal, or adult-T-suppressant treated individuals. This suggests a distinctive pattern of gene regulation in juvenile-GnRHa- treated individuals, highlighting unique aspects of the treatment’s impact on SSC gene expression. Furthermore, while AMH expression was traditionally assumed to be exclusive to Sertoli cells (*66*), our findings indicate a broader expression across pubertal tissues, encompassing Sertoli, Leydig, and SPG cells. The nuanced pattern of AMH inhibition in a cell-type-specific manner during puberty sheds light on a previously misunderstood source of AMH in serum.

In conclusion, the clinical specimens, data, and research strategy presented in this study provide unique opportunity to explore the impact of PB on testicular reproductive health.

## Methods

### Approval of Study and Patient Informed Consent

The Mayo Clinic Pediatric Testicular Biobank for Fertility Preservation operates on an opt-in basis, requiring written informed consent approved by the Mayo Institutional Review Board. Eligible donors, aged 0-17, must have parental consent. Participants consent to sample and data use, allowing access to questionnaires and medical records. Biopsies are collected for the clinical fertility preservation biobank and donate 20 % tissue for research. Recruitment primarily targets patients scheduled for pre-surgical appointments in the Department of Pediatric Urology at Mayo Clinic. The protocol was initiated in 2015 and first enrollment in 2016 at Mayo Clinic Rochester campus. The protocol grants access to stored clinical samples and deidentified data sharing with researchers. The consent document includes checkboxes for participants to indicate posthumous specimen designation. Collection of pediatric patient data and tissue adheres to Mayo Clinic Institutional Review Board guidelines, with parental consent obtained. Patients can withdraw consent at any time. Data including medical history, hormonal data, genetic data, and TESE outcomes were collected from electronic medical records. Prior-surgery, all donors were tested for Hepatitis B, Hepatitis C and HIV and stored in designated spaces. Biopsy tissue can be distributed to academic institutions working on fertility preservation research. Clinical information is summarized in **Table 1**.

### Surgical Procedure and Histological Analysis of Testicular Biopsy Specimen

Surgical procedures were conducted exclusively in the Department of Pediatric Urology at Mayo Clinic. All pediatric testicular biopsy specimens underwent systematic surgical procedures performed under general anesthesia. Following scrototomy and tunica vaginalis opening, the albuginea was incised, and the testicular pulp was expressed and dissected. Fragments were obtained bilaterally or unilaterally at the upper, middle, and lower poles of each testicle, avoiding the rete testis area. After transport to the laboratory, one fragment is sent to Mayo Clinic IVF Clinic for mechanical preparation and observation under optical microscopy for spermatozoa. Another fragment was sent to Anatomic Pathology, fixed, and stained with Hematoxylin and Eosin. Scanning was conducted using the Aperio GT450 whole slide image scanner (Leica Biosystems) in the clinical lab, and images were saved as SVS files. The digitized slide data were analyzed using QuPath software (version 0.4.3), as demonstrated in **Figure 2**, to assess seminiferous tubules.

### Testicular Specimen Processing for Single Cell Analysis

Fresh testicular specimen from the juvenile transgender donor was transported from the operating room to the processing laboratory. The specimen was washed in HBSS and subjected to a two-step enzymatic digestion as described previously (*28*). The digestion was stopped by adding HBSS containing 10 % FBS and passed through 40µm filter to obtain the single cell suspension. The cell suspension was centrifuged at 300 g for 5 min and resuspended in MEM medium with 10 % FBS.

The dissociated testicular cells were resuspended in PBS and stained with trypan blue and counted using a hemocytometer. The volume of cell suspension was adjusted to 1000 cells/µL in PBS. Cells were then processed on a 10x Genomics Chromium Controller using the Chromium Next GEM Single Cell 3′ reagents kit v3.1 (Dual Index) following the manufacturer’s instructions. In brief, approximately 16,500 live cells were loaded onto the Chromium controller to recover 10,000 cells for library preparation and sequencing. Gel beads were prepared following manufacturer’s instructions. Subsequently, oil partitions of single-cell and oligo coated gel beads were captured and reverse transcription was performed, resulting in cDNA tagged with a cell barcode and unique molecular index (UMI). Next, GEMs were broken, and cDNA was amplified and quantified using an Agilent Bioanalyzer High Sensitivity chip (Agilent Technologies).

To prepare the final libraries, amplified cDNA was enzymatically fragmented, end-repaired, and polyA tagged. Fragments were then size selected using SPRIselect magnetic beads (Beckman Coulter). Next, Illumina sequencing adapters were ligated to the size-selected fragments and cleaned up using SPRIselect magnetic beads (Beckman Coulter). Finally, sample indices were selected and amplified, followed by a double-sided size selection using SPRIselect magnetic beads (Beckman Coulter). Final library quality was assessed using an Agilent Bioanalyzer High Sensitivity chip. The libraries were then sequenced as paired end reads (PE150) on Novaseq platform (illumina). Raw sequencing reads were aligned to the human reference genome hg38, barcodes were processed and counted, and gene-barcode matrices were created using the *cellranger count* function of CellRanger software (v6.1.2).

### Bioinformatic Approaches

The *Seurat* (version 4.3.1.0) library was used to conduct scRNA-seq analysis in R (version 4.3.2). Due to the nature of meitogenetic process and to avoid data loss, we adopted a strategy wherein data was filtered for reads count and ribosome content. Unlike other tissues, cytoplasmic and nuclear compaction is known during end stages of spermatogenic differentiation (*67*) hence quality control filtering by mitochondrial content was excluded to avoid excessive loss of data. Following filtering we performed read count normalization using the *NormalizeData* function in *Seurat* with parameters of *LogNormalize* and scale factor of 10000 followed by variable feature identification using the *VariableFeatures* function. After filtering, count normalization and variable feature identification was conducted on each individual sample, *FindIntegrationAnchors* was run to scale and integrate the datasets. Subsequently, *IntegrateData* using canonical correlation analysis was run using these anchors. Finally, dimensionality reduction and data visualization of the integrated assay was conducted with *RunPCA* and *FindNeighbors* with 30 dimensions, *FindClusters* with resolution of 0.5, and *RunUMAP* with 30 dimensions. From this, 24 nearest neighbors were created. Clusters Unknown 1 and Sertoli 3 highly enriching for mitochondria were discarded and rest of the clusters were analyzed (**Supplementary Figure S1**). Individual prepubertal, pubertal, young adult, and old adult scRNAseq data from prior studies were labeled with meta-data on age and pubertal status prior to integration. The RNA assay and *DotPlot* on specific genes was subsequently used to do cell type identification. Cell-type identifications were subsequently stored in the integrated object’s metadata.

To ensure *Seurat’s* integration function was not forcing cell types to align with each other, we regressed out the effect of each patient batch and ran the standard *Seurat* pipeline to generate a UMAP. While most cell types aligned with each other regardless of batch, Unknown 1, Sertoli 4, Leydig 1, and Leydig 2 populations of the PB-patient were the only cell populations disconnected from the remaining patients (**Supplementary Figure S1B**). Upon analysis of markers expressed by these populations, Unknown 1 and Sertoli 4 of the PB-patient expressed only HBA1, HBA2, and mitochondrial genes (**Supplementary Figure S1A**). Therefore, these cells were excluded from analysis. Sertoli 3 was also excluded from analysis due to high mitochondrial gene expression in the RNA assay. Leydig 1 and Leydig 2 remained in analysis as the PB-patient still expressed standard Leydig markers.

Using the integrated assay, *DotPlot* was used to compare the spermatogenic and Sertoli cell markers across different age groups and treatments. UMAPs of spermatogenic and Sertoli lineages were developed by extracting lineage indices, creating a subsetted *Seurat* object, and running *RunPCA*, *FindNeighbors*, *FindClusters*, and *RunUMAP*. Custom functions were used to find proportions of cells within the spermatogenic and Sertoli lineages as well as within all cell types. Once proportions were found, we used the convex hull method in the *ggplot2* library to generate trend lines for normal patients and the *ggstar* library to label the T-suppressant/PB-treated patients. The *TernaryPlot* library was used to plot the relative proportions of Leydig, Sertoli, and spermatogenic lineages. Differential gene expression analysis was performed using the Wilcoxon rank-sum test and p-values were converted to q-values using the *qvalue* library for identification of pertinent gene.

To determine the expression of previously mentioned pubertal genes in individual clusters, *AggregateData* in the *tidyverse* library and grouping by object metadata. RNA assay was used to identify all pubertal genes detected and not limit based on the 2000 anchor features in the integrated assay. A limitation of this approach is that differences identified could be a result of batch-effects across samples. To further understand the age-based differences in expression of pubertal genes, individual Sertoli, Leydig, and spermatogenic cell-types were subsetted and *AggregateData* was performed sample meta-data was conducted.

### Machine Learning Approaches

We employed a generalized linear model binomial regression in *R* to generate a model that could use cell type proportions to: 1) Distinguish between incompetent (prepubertal) and competent (young adult) patients. 2) Give us an understanding of where remaining patients stand on the pubertal scale. 5 prepubertal patients ages 13 months (2), 2 years old (1), and 7 years old (2) and 5 young adult patients ages 21, 22 (n = 2), 24, and 25 were used to train this model. Pubertal, oldage and drug-treated patients were excluded from training. The resulting binomial generalized linear model (GLM) was applied to all datasets.

Additionally, we trained one random forest model for each cell type based on the previously mentioned incompetent and competent sample cells. The training dataset was a balanced dataset of incompetent and competent cells randomly selecting approximately 70 % of each class. The remaining 30 % of cells were left as a validation dataset. We only generated models for SPG 1, SPG 2, Sertoli 2, Sertoli 4, Sertoli 5, Leydig 1, Leydig 2, and Endothelial 1 since these clusters had at least 100 cells per class in the validation dataset, ensuring sufficient training and evaluation of the model. The h2o package was used to generate and optimize the model by iterating combinations of the following hyperparameters: L1 regularization of 1E-3, 2E-3, 5E-3, and 1E-2, L2 regularization of 1E-3, 2E-3, 5E-3, 1E-2, input dropout ratio of 0.1, 0.2, and 0.3, hidden dropout ratios of 0.2, 0.3, 0.4, and mini-batch size of 16, 32, and 64. The hyperparameter combination with highest validation accuracy was chosen using the h2o.confusionmatrix function. The default threshold to distinguish between classes was determined using the F1 score. The resulting models were applied to all datasets to generate prediction scores.

### Statistical analysis

Statistical methods for each analysis are detailed in the corresponding figure legends.

### Data availability

The accession number for public sequencing data analyzed are: GSE112013 (3 adult, 2x technical replicate) (*28*); GSE120506 (2 infant, 2x technical replicate) (*28*); GSE182786 (4 young and 5 old post-pubertal) (*30*); GSE196819 (3 pre-pubertal) (*30*)); GSE161617 (1 infant (2x technical replicate)(*32*); GSE134144 (1 pre-pubertal, 3 pubertal, and 2 adult puberty blocker, 2x technical replicate)(*10*).

## Acknowledgements

GK received Yamagiwa-Yoshida Memorial International Cancer (YY) study grants from UICC. V.M. received Mayo-Summer Undergraduate Fellowship. N.K. was supported by Mayo-NCI SPORE for Breast Cancer [CA116201-12CEP] and Ovarian Cancer [CA136393-11CEP] research. We specially thank research coordinators Brandi Johnson and Stephanie Hafner for data collection and Tiffany Mainella for their help with preparation of this manuscript. The authors dedicate this study in memory of Dr. Connie J. Eaves, a beloved mentor and pioneer in the field of stem cell research. Dr. Eaves had a profound impact on the careers of authors NK and DK.

## Author Contribution

CG, AC, and YZ contributed to the collection of clinical specimen and data. JG and SS provided assistance with histology analysis. GK, MA, and MR generated transgender scRNA data. VM, GK and NK performed the data analysis, prepared the figures and drafted the manuscript. DK contributed to data analysis and provided critical review of the manuscript. All authors contributed to the draft. NK funded and supervised the study.

## Supplementary Data

**Supplementary Table S1: *Differential expression analysis of prepubertal and adult spermatid 2 cells*.**

**Supplementary Table S2: *Differential expression analysis of prepubertal and adult spermatid 3 cells*.**

**Supplementary Table S3:** List of pubertal genes with non-zero expression in Myoid, Sertoli, Leydig, Macrophage, and Endothelial clusters

**Supplementary Table S4: *Evaluation of machine learning model accuracy.*** The area-under-the receiver operator curve (AUC) using training and test datasets consisting of prepubertal and adult samples.

*Supplementary Table S5:* List of relative and scaled importance as well as percentage contribution for all genes in each machine-learning model

**Supplementary Figure S1: *Nontraditional quality control filtering and standard cell type identification.* A)** Dot plot of mitochondrial genes per cell cluster used to exclude certain clusters with high mitochondrial content. **B)** UMAPs split by age ranges and cell types that result from regressing out sample-to-sample scRNAseq processing variation, showing cell-types are common between each batch. **C)** Dot plot showing the expression of standard cell-type specific genes used in cell-cluster identification within the PB-treated patient.

**Supplementary Figure S2: *Cell type identification using cells from all patients.* A)** Dot plot of spermatogenic lineage cell markers. **B)** Dot plot of Sertoli and Leydig lineage cell markers. **C)** Dot plot of endothelial and myoid lineage cell markers. **D)** Dot plot of macrophage cell markers.

**Supplementary Figure S3: *Cell type identification using cells from all patients.*** Trends in the relative cell proportions with normal development highlighted in honeydew blue. The red star represents the PB-treated patient, whereas blue and yellow stars respectively represent the 26-year-old and 50-year-old T-suppressant-treated patients.

**Supplementary Figure S4: *Impact of puberty blocker and aging on all cell types.*** Heatmap representing pubertal genes found in RNA assay versus pseudo-bulked *Seurat* object by cell-type.

**Supplementary Figure S5: *Impact of puberty blocker in sample-to-sample in Sertoli, Leydig, and early spermatogonia.*** Heat map pubertal genes found in integrated assay versus pseudo-bulked *Seurat* object by sample. Each heatmap in panel represents subsetted cells from Sertoli, Leydig, and spermatogenic clusters.

**Supplementary Figure S6: *Upregulated and downregulated genes in GnRHa versus pubertal patients for each cell type.*** Volcano plots comparing GnRHa and Pubertal samples for each cell type using q-value, excluding cells that had no differentially expressed genes above the q-value threshold of 0.001 and absolute log- fold change of 0.5.

**Supplementary Figure S7: *Changes in standard Sertoli marker expression across sample categories.*** Dot plot depicting relative expression of standard Sertoli lineage cell markers from existing literature between developmental categories and treatment received, if applicable.

**Supplementary Figure S8: *Importance of topmost features in machine learning models*.**

**Supplementary Figure S9: *Machine learning prediction score distributions of cells within individual clusters.*** Prediction score distributions in cell-types with enough Age 0-7 and Age 21-25 cells to separate into training and validation. For Age 0-7 and Age 21- 25 score distributions, both training and validation indices are included.

## Notes

### Competing Interest Statement

The authors have declared no competing interest.

